# Reproductive tissues-specific meta-QTLs and candidate genes for development of heat-tolerant rice cultivars

**DOI:** 10.1101/2020.05.02.073429

**Authors:** Qasim Raza, Awais Riaz, Khurram Bashir, Muhammad Sabar

## Abstract

Rice holds the key to future food security. In rice-growing areas, temperature has already reached an optimum level for growth, hence, any further increase due to global climate change could significantly reduce rice yield. Several mapping studies have identified a plethora of reproductive tissue-specific and heat stress associated inconsistent quantitative trait loci (QTL), which could be exploited for improvement of heat tolerance. In this study, we performed a meta-analysis on previously reported QTLs and identified 35 most consistent meta-QTLs (MQTLs) across diverse genetic backgrounds and environments. Genetic and physical intervals of nearly 66% MQTLs were narrower than 5 cM and 2 Mb respectively, indicating hotspot genomic regions for heat tolerance. Comparative analyses of MQTLs underlying genes with microarray and RNA-seq based transcriptomic data sets revealed a core set of 45 heat-responsive genes, among which 24 were reproductive tissue-specific and have not been studied in detail before. Remarkably, all these genes corresponded to various stress associated functions, ranging from abiotic stress sensing to regulating plant stress responses, and included heat-shock genes (*OsBiP2, OsMed37_1*), transcription factors (*OsNAS3, OsTEF1, OsWRKY10, OsWRKY21*), transmembrane transporters (*OsAAP7A, OsAMT2;1*), sugar metabolizing (*OsSUS4*, α*-Gal III*) and abiotic stress (*OsRCI2-7, SRWD1*) genes. Functional data evidences from *Arabidopsis* heat-shock genes also suggest that *OsBIP2* may be associated with thermotolerance of pollen tubes under heat stress conditions. Furthermore, promoters of identified genes were enriched with heat, dehydration, pollen and sugar responsive cis-acting regulatory elements, proposing a common regulatory mechanism might exist in rice for mitigsating reproductive stage heat stress. These findings strongly support our results and provide new candidate genes for fast-track development of heat-tolerant rice cultivars.

**Key Message:** By integrating genetics and genomics data, reproductive tissues-specific and heat stress responsive 35 meta-QTLs and 45 candidate genes were identified, which could be exploited through marker-assisted breeding for fast-track development of heat-tolerant rice cultivars.

## Introduction

Rice (*Oryza sativa* L.) holds the key to future food security, as more than 50% of the world’s population solely depends on it for meeting their daily calorie requirements (Confalonieri et al. 2009). Rice has its origin in tropical and subtropical areas and considered as a thermophilic species (Yin et al. 1996). During the 21^st^ century, an increase of 1.5–4.5 °C in air temperature is predicted due to global climate change (Peraudeau et al. 2015). Rice is already growing in those agricultural areas where temperature has reached an optimum level (33/25 °C) for growth (Yin et al. 1996) and a further increase in day or night temperatures during heat-sensitive growth stages could significantly reduce rice yield (Krishnan et al. 2011). Like other crop plants, rice is most susceptible to reproductive stage heat stress (Satake and Yoshida 1978; Prasad et al. 2017). High temperature at this stage causes irreversible damage by retarding anther dehiscence, pollen fertility, germination on stigma, pollen tube elongation in pollinated pistils, spikelet fertility and grain filling (Jagadish et al. 2007; Jagadish et al. 2010b; Maruyama et al. 2013; Shah et al. 2011; Zhang et al. 2018). Reproductive stage heat tolerance of rice could be improved by elucidating complex molecular-genetic regulatory mechanisms underlying heat tolerance and genetic exploitation of available heat-tolerant sources.

In-depth understating of stress response mechanisms is extremely important for successful development of stress-tolerant crops. Survival under heat stress is dependent upon plants ability to perceive stress stimulus, generate and transmit stress signal and initiate appropriate physiological, biochemical and molecular changes (Shanmugavadivel et al. 2019, Bashir et al. 2019). Heat stress causes overproduction of reactive oxygen species (ROS), underproduction of protective enzymes and proteins and degradation of the plasma membrane, which in turn leads to cell death. Plants respond to heat stress by activating their heat-responsive genes. These heat-inducible genes activate antioxidant enzymes and osmoprotectants to detoxify ROS, reactivate essential enzymes and structural proteins and re-establish the cellular homeostasis (Hasanuzzaman et al. 2013). Heat-shock protein (*HSP*s)-encoding genes are master players for heat tolerance in plants (Yadav et al. 2020). During heat stress, the expression of *HSPs* is upregulated, which in turn activates the chaperones network to protect intracellular proteins from denaturation and preserves their stability by protein folding (Baniwal et al. 2004). These *HSP*s encompass some conserved cis-acting regulatory elements (CREs) in their promoter regions which trigger their transcription in response to heat stress (Nover et al. 2001).

To date, a plethora of QTLs for reproductive stage heat tolerance has been identified in different mapping populations under controlled and field environments (Buu et al. 2014; Cao et al. 2003; Cheng et al. 2012; Jagadish et al. 2010a; Poli et al. 2013; Qingquan et al. 2008; Shanmugavadivel et al. 2017; Udawela et al. 2018; Xiao et al. 2011a; Xiao et al. 2011b; Ye et al. 2012; Ye et al. 2015a; Zhang et al. 2009; Zhang T et al. 2008; Zhao et al. 2016; Zhao et al. 2006; Zhu et al. 2017). However, inconsistent QTL information is provided in these studies because different mapping populations, experimental designs, statistical analyses and environments were involved. During recent years, meta-analysis approach has become a mainstream practice for integrating the QTL information data obtained from multiple independent mapping experiments and identification of most consistent meta-QTLs (MQTLs) across diverse environments. This approach has been applied in several economically important food crops, including maize, wheat, barley and rice (Martinez et al. 2016; Darzi-Ramandi et al. 2017; Khahani et al. 2019; Raza et al. 2019). These studies provided a proof of concept for identification and prioritization of trait linked MQTLs/candidate genes and offered new biological data for the development of stress tolerant rice cultivars.

In this study, we executed a meta-analysis on previously reported reproductive tissue associated QTLs to identify most consistent MQTLs across different experiments and environments. Then, we integrated MQTLs and transcriptomic data to prioritize heat-responsive candidate genes. We also demonstrated a potential relationship between thermotolerance of pollen tubes and *OsBIP2*. Finally, we performed an in-silico promoter analysis on heat-responsive genes to find common CREs which trigger a heat stress response. These findings offer new candidate genes for the development of heat-tolerant rice cultivars.

## Material and Methods

### Data retrieval

Recent literature published in journal articles regarding mapping of reproductive stage heat stress associated QTLs in rice were retrieved by the search of Google Scholar. Several independent studies have mapped 163 reproductive tissues associated QTLs with proper genetic map information onto the rice chromosomes (**Table 1**). Background data (chromosomal location and position, confidence interval range, log of odds (LOD) and phenotypic variance etc.) of all these QTLs were mined from the original studies (see **Table 1**). Missing confidence interval values were estimated using equations proposed by Darvasi and Soller (1997). If LOD and phenotypic variance values were missing from the original study, we assumed these as 2.5 and 10%, respectively (Khahani et al. 2019).

**Table 1.** Reproductive stage QTL mapping studies in rice

| **Study** | **Parents** | **Population** | **Size** | **QTLs\*** |
| --- | --- | --- | --- | --- |
| Cao et al. (2003) | IR64 × Azucena | DH | 109 | 6 |
| Zhao et al. (2006) | USSR5 × Guangjie9// USSR5 | BC | 7, 123 | 5 |
| Qingquan et al. (2008) | T219 × T226 | RIL | 202 | 7 |
| Zhang et al. (2008) | Zhongyouzao8 × Toyonishiki | RIL | 168 | 5 |
| Zhang et al. (2009) | 996 × 4628 | RIL | 279 | 2 |
| Jagadish et al. (2010a) | Bala × Azucena | RIL | 181 | 16 |
| Xiao et al. (2011a) | 996 × 4628 | RIL | 286 | 2 |
| Xiao et al. (2011b) | 996 × 4628 | RIL | 286 | 2 |
| Ye et al. (2012) | IR64 × N22 | BC, RIL | 188, 158 | 15 |
| Cheng et al. (2012) | Xiushui9 × IR2061 | BC | 240 | 2 |
| Poli et al. (2013) | IR64 × NH219 | RIL | 70 | 2 |
| Buu et al. (2014) | OM5930 × N22 | BC | 310 | 11 |
| Ye et al. (2015a) | IR64 × N22 | RIL | 96 | 11 |
|  | IR64 × Giza178 | RIL | 86 |  |
|  | IR64//Milyang23 × Giza178 | RIL | 166 |  |
| Zhao et al. (2016) | Sasanishiki × Habataki | CSSL | 37 | 7 |
| Shanmugavadivel et al. (2017) | N22 × IR64 | RIL | 272 | 5 |
| Zhu et al. (2017) | Sasanishiki × Habataki | CSSL | 39 | 12 |
| Udawela et al. (2018) | Bg 90 – 2 × SH 527 | BC | 400 | 53 |
|  | Mengguandamagu × SH 527 |  |  |  |

\*, number of reproductive tissues-specific QTLs.

### Meta-QTL analysis

Before employing MQTL analysis, a consensus map was developed by projecting individual genetic maps onto the Cornoll SSR 2001 rice genetic map (Temnykh et al. 2001). Only those QTL clusters were considered for final MQTL analysis which showed overlapping positions onto respective chromosomes during at least two independent mapping studies. BioMercator V4.2 (Sosnowski et al. 2012) was employed to detect hotspot MQTL regions. Based on the Akaike information criterion (AIC), BioMercator proposes five QTL models. We choose the lowest AIC value containing model because it loses less information and considered the best model (Akaike 1998).

### Identification of candidate genes

Reproductive tissues associated heat-responsive genes were identified by following two steps. In the first step, physical locations of markers flanking the confidence intervals of identified rice MQTLs (rMQTL’s) were obtained from Gramene Annotated Rice Genome 2009. If the physical position of a marker was absent, then we used position of adjacent genetic markers. All genes positioned within physical intervals of MQTL regions were extracted as batch download from Rice Annotation Project Database (Sakai et al. 2013). In the second step, non-redundant genes obtained from the first step were compared with heat-responsive genes of two previously published genome-wide transcriptome studies conducted during rice reproductive development. (Zhang et al. 2012) investigated time-course gene expression profile in young panicles of a heat-tolerant rice variety 996 and identified 2449 differentially expressed genes (DEGs). Similarly, (González-Schain et al. 2016) performed a transcriptome study between heat-tolerant and heat-sensitive rice genotypes under heat-stress conditions during anthesis and identified 630 DEGs. Common genes between the two above mentioned studies and current work were identified and considered as a core set of reproductive tissues associated heat-responsive candidate genes. Gene Ontology (GO) and sub-location information of all genes were collected from RiceNetDB (Liu et al. 2013) and Wolf PSORT (Horton et al. 2007) databases, respectively. Gene evidence network was generated with Knetminer, a publically available database (Hassani-Pak et al. 2016), by exploring O*sBIP2* (*Os03g0710500*) connection with heat tolerance using default parameters.

### Promoter analysis

The 1.5 kb nucleotide sequences upstream from the translational start site (ATG) of all genes were downloaded from Phytozome database and uploaded in PLACE database (Higo et al. 1999) for prediction of common cis-acting regulatory elements (CREs) and their putative functions. For simplicity, only those CREs were considered which occurred at least once in > 80% of the total candidate genes.

## Results

### Reproductive tissues-specific MQTLs in rice

Background information on reproductive tissue-specific and heat-stress associated QTLs were retrieved from 17 independent reports published since 2003, corresponding to 12 distinct and 3 redundant mapping populations (**Table 1**). Several vegetative tissues-specific and critical information lacking QTL studies published on this topic were not included in current analysis (Bahuguna et al. 2015; Kilasi et al. 2018; Singh et al. 2017). Seventeen studies used for current MQTL analysis covered diverse population types and sizes. A total of 163 reproductive tissue-specific QTLs were reported in these studies and were unevenly distributed on twelve rice chromosomes. During meta-analysis, 122 initial QTLs (approx. 75%) showed overlapping positions between at least two independent studies and were included in the final analysis.

Based on the lowest AIC value, 35 rMQTL’s were detected on twelve rice chromosomes. A maximum number of rMQTL’s were located on 2^nd^, 3^rd^, 4^th^ and 11^th^ chromosomes (four on each), whereas minimum was located on 7^th^ and 12^th^ chromosomes (one each) (**Fig. 1**). Each rMQTL harboured at least two initial QTLs from two independent studies. The highest number of initial QTLs were encompassed by rMQTL10.3 (9), followed by rMQTL6.1 (8), rMQTL2.4 (6) and rMQTL10.1 (6) indicating the most stable MQTLs across diverse experimental conditions. After MQTL analysis, significant reductions in confidence intervals (CI’s) of initial QTLs were observed. The 95% CI’s of recognized rMQTL’s ranged from 0.30 – 13.77 cM with a median value of 3.03 cM (**Table 2**). Likewise, the physical intervals of identified rMQTL’s were also reduced and varied from 0.04 – 5.85 Mb with a median value of 1.52 Mb. Interestingly, 23 rMQTL’s (approx. 66%) exhibited less than 5 cM CI’s and narrower than 2 Mb physical intervals, suggesting these are important heat-responsive genomic regions during reproductive development in rice. Details of all identified rMQTL’s are provided in **Table 2**.

**Table 2.** Detailed information of reproductive tissue-specific meta-QTLs.

| Sr No. | MQTL | Chr | CI Range (cM) | Position (cM) | Left marker |  |  | Right marker |  |  | No. of initial QTLs | CI (95%) (cM) | Physical Interval (bp) | Number of genes |
| --- | --- | --- | --- | --- | --- | --- | --- | --- | --- | --- | --- | --- | --- | --- |
|  |  |  |  |  | Name | Position (cM) | Position (bp) | Name | Position (cM) | Position (bp) |  |  |  |  |
| 1 | rMQTL1.1 | 1 | 115.16 - 127.87 | 121.52 | RG345 | 115.2 | 27118807 | RM403 | 131.2 | 29384826 | 3 | 12.71 | 2.27 | 319 |
| 2 | rMQTL1.2 | 1 | 154.31 - 158.32 | 156.31 | RM486 | 153.5 | 34955554 | RM315 | 165.3 | 36734272 | 3 | 4.01 | 1.78 | 247 |
| 3 | rMQTL1.3 | 1 | 205.63 - 207.68 | 206.66 | RG331 | 194.1 | 41540722 | RM1067 | 208.8 | 42923549 | 5 | 2.05 | 1.38 | 220 |
| 4 | rMQTL2.1 | 2 | 7.89 - 16.43 | 12.16 | RM110 | 6.9 | 1326947 | RM233 | 16.3 | 2070184 | 2 | 8.54 | 0.74 | 115 |
| 5 | rMQTL2.2 | 2 | 70.53 - 84.3 | 77.41 | RM262 | 70.2 | 20794972 | RG157 | 90.3 | 19866086 | 3 | 13.77 | 0.93 | 75 |
| 6 | rMQTL2.3 | 2 | 142.82 - 150.95 | 146.89 | RM599 | 139.6 | 27109220 | RM450 | 150.8 | 28628356 | 5 | 8.13 | 1.52 | 171 |
| 7 | rMQTL2.4 | 2 | 186.17 - 186.99 | 186.58 | RM208* | 186.4 | 35135783 | RM482 | 187.5 | 35272560 | 6 | 0.82 | 0.14 | 26 |
| 8 | rMQTL3.1 | 3 | 1.45 - 1.75 | 1.60 | RM60 | 0.0 | 105852 | RG104 | 3.5 | 485333 | 2 | 0.30 | 0.38 | 69 |
| 9 | rMQTL3.2 | 3 | 99.54 - 100.95 | 100.25 | RM3280 | 90.4 | 10903514 | RM338 | 108.4 | 13221665 | 3 | 1.41 | 2.32 | 296 |
| 10 | rMQTL3.3 | 3 | 162.96 - 165.32 | 164.14 | CDO337 | 161.1 | 28231692 | RM1350 | 165.5 | 28676570 | 4 | 2.36 | 0.44 | 54 |
| 11 | rMQTL3.4 | 3 | 202.49 - 204.17 | 203.33 | RM468 | 202.3 | 32674852 | RM422 | 205.4 | 33709541 | 2 | 1.68 | 1.03 | 188 |
| 12 | rMQTL4.1 | 4 | 7.97 - 9.36 | 8.66 | RM16251** | - | 48875 | RM537 | 15.0 | 176875 | 5 | 1.39 | 0.13 | 8 |
| 13 | rMQTL4.2 | 4 | 66.4 - 70.36 | 68.38 | RM1155 | 58.9 | 20344188 | RM564 | 73.1 | 18587434 | 3 | 3.96 | 1.76 | 184 |
| 14 | rMQTL4.3 | 4 | 80.47 - 82.88 | 81.67 | RZ565 | 80.1 | 21451046 | RZ675 | 94.4 | 24165408 | 2 | 2.41 | 2.71 | 416 |
| 15 | rMQTL4.4 | 4 | 111.85 - 118.2 | 115.03 | RM241 | 106.2 | 26857374 | RM317 | 118.3 | 29061127 | 5 | 6.35 | 2.20 | 276 |
| 16 | rMQTL5.1 | 5 | 8.79 - 15.37 | 12.08 | RM6300 | 6.5 | 600257 | RM413 | 26.7 | 2212825 | 3 | 6.58 | 1.61 | 208 |
| 17 | rMQTL5.2 | 5 | 96.92 - 99.97 | 98.44 | RM305 | 96.9 | 20944257 | RM3790 | 100.4 | 26195611 | 2 | 3.05 | 5.25 | 693 |
| 18 | rMQTL5.3 | 5 | 118.36 - 121.39 | 119.88 | RZ70 | 112.0 | 24261900 | RM26 | 122.7 | 27342124 | 2 | 3.03 | 3.08 | 184 |
| 19 | rMQTL6.1 | 6 | 6.96 - 7.7 | 7.33 | RM589 | 3.2 | 1380866 | RM587 | 10.7 | 2292076 | 8 | 0.74 | 0.91 | 161 |
| 20 | rMQTL6.2 | 6 | 25.28 - 32.12 | 28.70 | RM204 | 25.1 | 3168314 | RM111 | 35.3 | 5096867 | 2 | 6.84 | 1.93 | 285 |
| 21 | rMQTL6.3 | 6 | 74.39 - 76.38 | 75.38 | RG648 | 74.3 | 19337571 | RM465B* | 75.5 | 16750851 | 3 | 1.99 | 2.59 | 136 |
| 22 | rMQTL7.1 | 7 | 105.91 - 112.56 | 109.24 | RZ978 | 103.3 | 28411532 | RG351 | 115.3 | 29467498 | 2 | 6.65 | 1.06 | 158 |
| 23 | rMQTL8.1 | 8 | 55.51 - 58.29 | 56.90 | RM3572 | 54.5 | 3927309 | RM344 | 61.9 | 9778963 | 4 | 2.78 | 5.85 | 468 |
| 24 | rMQTL8.2 | 8 | 99.16 - 108.46 | 103.81 | RM531 | 90.3 | 22469338 | RZ66 | 114.9 | 25592993 | 2 | 9.30 | 3.12 | 368 |
| 25 | rMQTL9.1 | 9 | 2.72 - 3.87 | 3.30 | CDO590 | 1.8 | 3759711 | RM5799 | 5.5 | 3802909 | 3 | 1.15 | 0.04 | 3 |
| 26 | rMQTL9.2 | 9 | 58.05 - 63.89 | 60.97 | RM434 | 57.7 | 15662573 | RM410 | 64.4 | 17643295 | 2 | 5.84 | 1.98 | 245 |
| 27 | rMQTL9.3 | 9 | 84.65 - 89.36 | 87.01 | RM160 | 83.9 | 19788222 | RG667 | 90.7 | 20482185 | 3 | 4.71 | 0.69 | 112 |
| 28 | rMQTL10.1 | 10 | 16.74 - 18.13 | 17.44 | RM7217** | - | 4278021 | RM216 | 17.6 | 4986996 | 6 | 1.39 | 0.71 | 44 |
| 29 | rMQTL10.2 | 10 | 68.14 - 76.6 | 72.37 | RZ625 | 58.3 | 16619676 | G2155 | 82.5 | 19823299 | 2 | 8.46 | 3.20 | 411 |
| 30 | rMQTL10.3 | 10 | 109.92 - 110.82 | 110.37 | RM333 | 110.4 | 22372009 | RM496 | 113.0 | 22430420 | 9 | 0.90 | 0.06 | 6 |
| 31 | rMQTL11.1 | 11 | 0.28 - 0.91 | 0.60 | RM4B | 0.0 | 932068 | RM558B | 8.2 | 1609092 | 4 | 0.63 | 0.68 | 86 |
| 32 | rMQTL11.2 | 11 | 35.7 - 40.3 | 38.00 | RM332 | 27.9 | 2840211 | RM552 | 40.6 | 4843207 | 3 | 4.60 | 2.00 | 195 |
| 33 | rMQTL11.3 | 11 | 76.83 - 78.68 | 77.75 | RM209 | 73.9 | 17808335 | RM457 | 83.0 | 19064994 | 2 | 1.85 | 1.26 | 73 |
| 34 | rMQTL11.4 | 11 | 101.38 - 108.72 | 105.05 | RM473E | 95.6 | 25457233 | RG1109* | 106.5 | 23155291 | 3 | 7.34 | 2.30 | 193 |
| 35 | rMQTL12.1 | 12 | 4.92 - 5.92 | 5.42 | RM20A | 3.2 | 970538 | RG574 | 13.2 | 1595325 | 4 | 1.00 | 0.62 | 92 |

\*, marker located within MQTL confidence interval range; \*\*, nearest possible marker from IRGSP genetic map 2005.

**Fig. 1.**
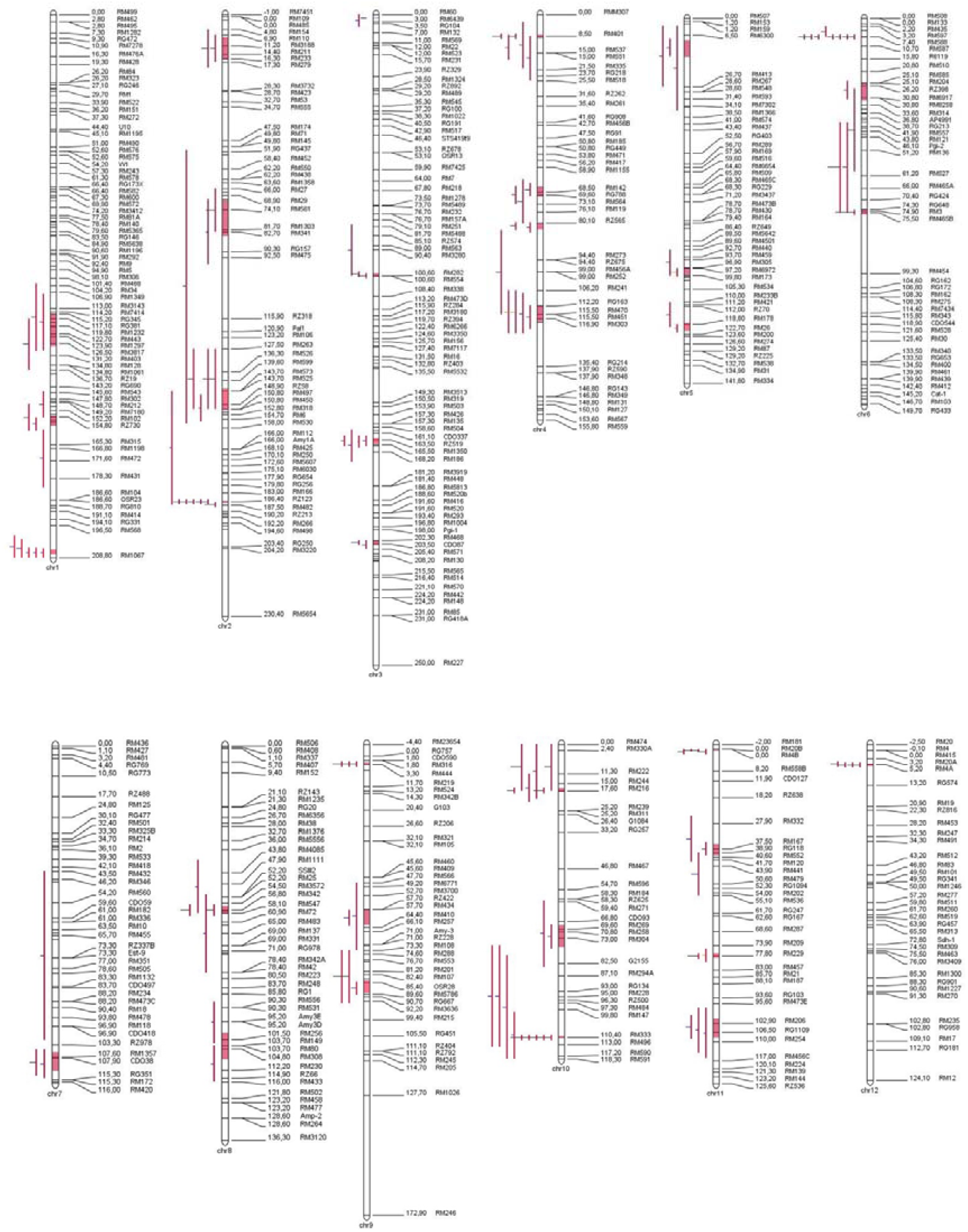
Graphical representation of meta-QTLs on rice chromosomes. Vertical bars with names at the bottom represent twelve rice chromosomes. Marker names and positions (cM) are shown on the left side of each chromosome. Positions and confidence intervals of original QTLs are presented as red lines on the right side of respective chromosomes. Whereas, red portions on chromosomes represent confidence intervals of identified meta-QTLs.

### Heat-responsive candidate genes within MQTL regions

All genes physically located within identified MQTL regions were considered as candidate genes and retrieved from RAP-DB as described in material and methods section. A total of 6785 non-redundant genes were underlying 35 rMQTL’s and their detailed information is provided in **Supplementary File-sheet 1**. Highest numbers of genes were encompassed by rMQTL5.2 (693), followed by rMQTL8.1 (468), rMQTL4.3 (416) and rMQTL10.2 (411) (**Table 2**). Whereas, rMQTL9.1, rMQTL10.3 and rMQTL4.1 harboured only 3, 6 and 8 genes, respectively. Overall, rMQTL’s with narrower physical intervals had a small number of genes and those with wider physical intervals had a large number of genes.

To identify a core set of reproductive tissues associated genes, we compared all 6785 genes identified in this study with heat-responsive DEGs of two independently published reports (Zhang et al. 2012; González-Schain et al. 2016). This comparison revealed a higher number of common genes between current work and Zhang et al. (2012) (439 genes), than with González-Schain et al. (2016) (126 genes) (**Supplementary File-sheets 2–4**). Difference in the number of common genes among three studies might be due to diverse genetic backgrounds (996, N22 and diverse populations), molecular techniques and specificity of expression patterns during different developmental stages. Interestingly, our analysis identified a core set of 45 most consistent heat-responsive genes which were common to all three studies (**Fig. 2A**). This core set included nine metabolic enzymes, seven heat shock proteins (HSPs), four transcription factors, three transporters, two reactive oxygen species (ROS) related, two ubiquitin-proteasome system (UPS) related and 18 other unclassified genes (**Supplementary File-sheet 5**). GO enrichment analysis revealed > 30% genes were involved in stress response ‘biological process’ and subcellular localization analysis exhibited predominant occurrence of > 70% genes in cytosol, nucleus and chloroplast (**Fig. 2B**). Remarkably, majority of these genes were differentially upregulated in reproductive tissues, especially ovary, embryo and endosperm, under normal field conditions as reported by Sato et al. (2011a) and (2011b) (**Fig. 3**). We noticed that 76% (34 out of 45) of our core set of genes were up-regulated, whereas 24% (11 out of 45) were down-regulated between heat-tolerant and sensitive rice genotypes in González-Schain et al. (2016) study. Interestingly, all *HSP*s were up-regulated in reproductive tissues of heat-tolerant rice cultivar N22, indicating high temperature-dependent activation of chaperones network. González-Schain et al. (2016) also identified a core set of 37 heat-responsive genes and we compared our core set of genes with their data set. Six genes were common between their and current work core sets, among which four were heat shock protein-encoding genes (*HSP*s), whereas 39 were unique. These unique genes can be exploited through marker-assisted selection (MAS) for development of heat-tolerant rice cultivars.

**Fig. 2.**
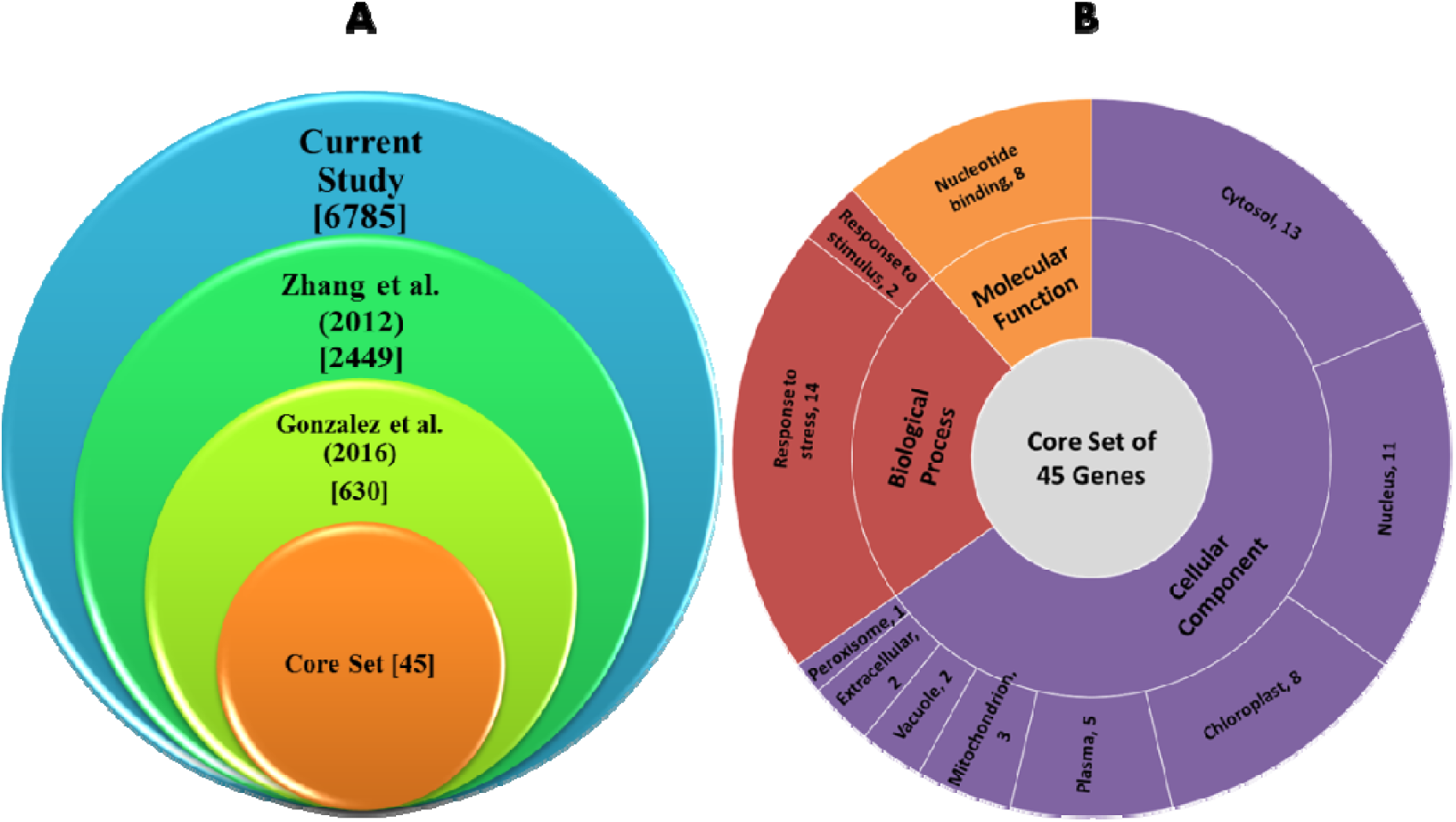
Comparison of genes among three data sets and go enrichment analysis of core set of heat-responsive genes. **(A)** Stacked Venn diagram showing comparison among three data sets for identification of common heat-responsive candidate genes. Numbers in square brackets indicate gene number in respective study **(B)** Sunburst showing GO enrichment analysis of core set of 45 heat-responsive genes. The outermost cluster represents different biological, cellular and molecular groups, whereas numbers at the end of each group indicate the number of individual genes falling in that group.

**Fig. 3.**
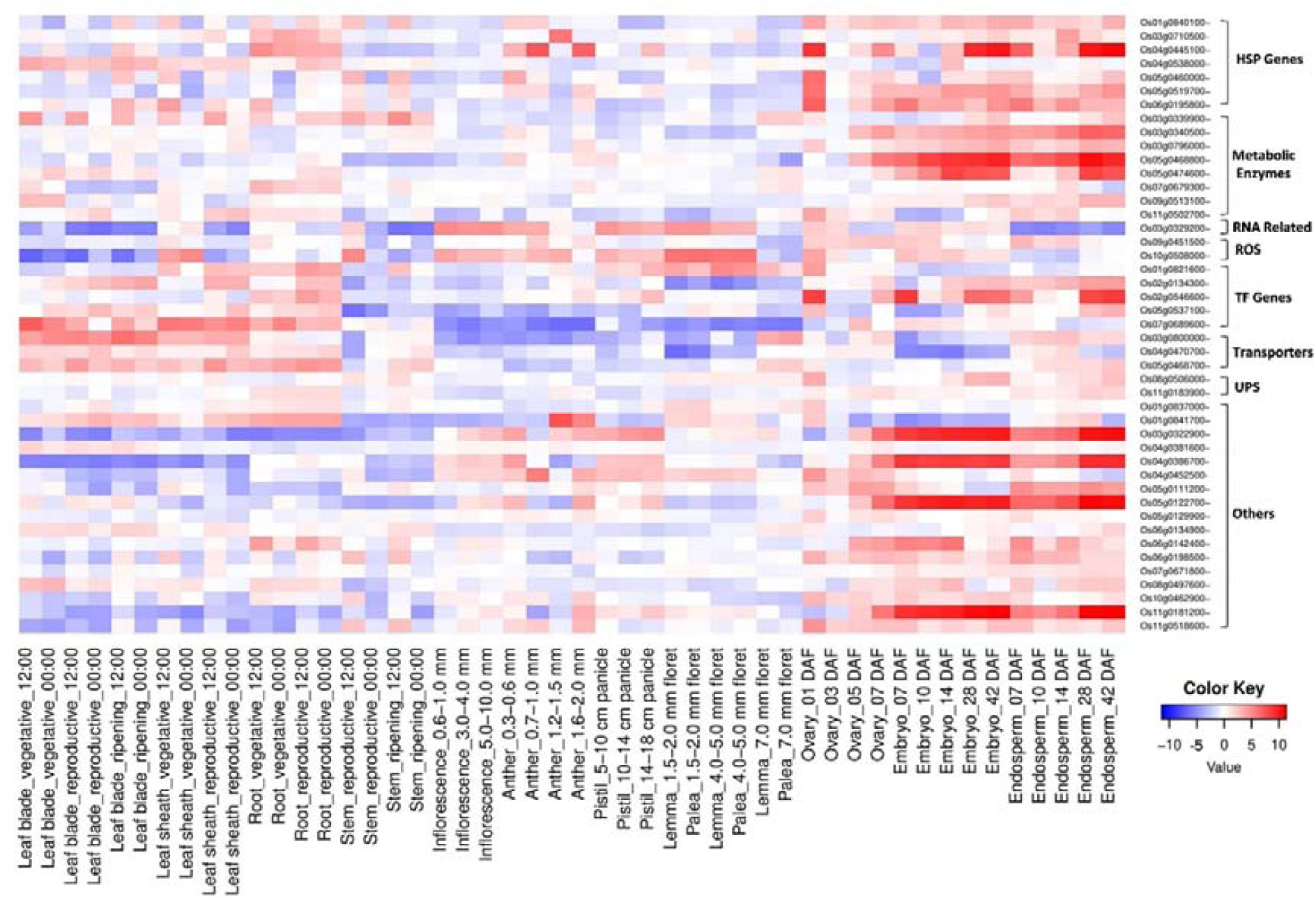
Expression patterns of core set of heat-responsive genes under natural field conditions. The normalized (Log_2_) expression data were mined from RiceXPro (http://ricexpro.dna.affrc.go.jp). Details of expression analysis and normalization were described by Sato et al. (2011a, 2011b).

We also assumed that some of our core set of heat-responsive genes might also contribute to heat response at the seedling stage. To investigate this likelihood, we again compared our core set of genes with another published study describing the differential expression of genes in heat-stressed seedlings (Mittal et al. 2012). Almost 47% of our core set of genes (21 out of 45) were also differentially expressed at seedling stage indicating a core set of basal heat-responsive genes in rice (**Supplementary File-sheet 5**) while remaining 53% (24 out of 45) appears to be reproductive tissue-specific genes (**Table 3**). Remarkably, all these genes corresponded to various stress associated functions, ranging from abiotic stress sensing to regulating plant physiological, biochemical and molecular responses, and included heat shock proteins, transcription factors, antioxidant enzymes, transmembrane transporters and sugar metabolism-related enzymes. These reproductive tissue-specific genes were differentially deregulated under heat stress conditions in both microarray and RNA-seq based transcriptome studies. Interestingly, one gene (*OsBiP2*; *Os03g0710500*) was highly expressed at all-time points (log_2_ FC > 2.5) and showed early heat response in both studies. Functional data evidences from *Arabidopsis* heat-shock genes suggested that *OsBIP2* might be associated with thermotolerance of pollen tubes under heat stress conditions (**Fig. 4**). We also observed comparable gene expression profiles in microarray and RNA-seq methods, with an exception at few points, suggesting candidate genes identified in current work are truthfully heat-responsive and directly or indirectly contribute to reproductive stage heat-tolerance. Enclosure of the majority of these genes in cross-reference studies further validates our results (**Table 3**).

**Table 3.** Reproductive tissue-specific heat-responsive genes in rice

| RAP (OS ID) | MSU (LOC ID) | Gene Name | Description | Log2FC<sup>1</sup> | Log2FC (20-min)<sup>2</sup> | Log2FC (60-min)<sup>2</sup> | Log2FC (2-h)<sup>2</sup> | Log2FC (4-h)<sup>2</sup> | Log2FC (8-h)<sup>2</sup> |
| --- | --- | --- | --- | --- | --- | --- | --- | --- | --- |
| Os01g0821600<sup>4</sup> | LOC_Os01g60640 | *OsWRKY21* | WRKY transcription factor 21. | -2.84 | 0.02 | -0.52 | -1.80 | -2.65 | -1.63 |
| Os01g0837000<sup>3</sup> | LOC_Os01g61990 | *OsNPR4* | Ankyrin repeat containing protein. | -1.53 | 2.17 | 1.78 | 1.01 | 1.45 | 1.29 |
| Os01g0840100<sup>3</sup> | LOC_Os01g62290 | *OsMed37_1* | Heat shock protein Hsp70 family protein. | 1.49 | 1.69 | 1.67 | 1.51 | 1.03 | 0.84 |
| Os01g0841700 | LOC_Os01g62430 | *OsERG1* | Similar to Elicitor-responsive protein 1. | -1.96 | 2.71 | 2.62 | 2.41 | 1.38 | 2.05 |
| Os02g0134300<sup>3</sup> | LOC_Os02g04160 | *OsTEF1* | Protein of unknown function DUF701. | 2.28 | 1.91 | 4.14 | 3.53 | 3.88 | 3.25 |
| Os03g0322900<sup>5</sup> | LOC_Os03g20680 | *OsLEA17* | Late embryogenesis abundant (LEA) protein. | 1.44 | 2.34 | 1.84 | 1.71 | 0.22 | 2.33 |
| Os03g0329200 | LOC_Os03g21160 | *OsC3H23* | Nucleotide-binding, alpha-beta domain protein. | -1.02 | -0.83 | -0.63 | -2.01 | -1.97 | -2.07 |
| Os03g0340500<sup>3</sup> | LOC_Os03g22120 | *OsSUS4* | Similar to Sucrose synthase. | 1.19 | 1.59 | 2.14 | 2.39 | 2.48 | 2.58 |
| **Os03g0710500<sup>3</sup>** | **LOC_Os03g50250** | ***OsBiP2*** | **Similar to Luminal binding protein.** | **4.16** | **6.35** | **7.14** | **4.68** | **2.67** | **2.67** |
| Os03g0800000 | LOC_Os03g58580 |  | Similar to Nitrate and chloride transporter. | -1.41 | -3.28 | -2.66 | -1.42 | -0.62 | -0.16 |
| Os04g0386700 | LOC_Os04g31710 |  | Similar to OSIGBa0148P16.6 protein. | 1.45 | 3.05 | 3.60 | 4.08 | 3.32 | 3.30 |
| Os04g0470700 | LOC_Os04g39489 | *OsAAP7A* | Amino acid transporter. | 1.05 | 3.31 | 3.69 | 2.58 | 0.85 | 1.01 |
| Os05g0111200 | LOC_Os05g02060 |  | Similar to Amino acid selective channel protein. | 1.07 | 1.58 | 2.32 | 2.06 | 1.71 | 0.65 |
| Os05g0122700<sup>3</sup> | LOC_Os05g03130 | *OsRCI2-7* | Similar to Low-temperature induced protein. | 2.13 | 1.64 | 2.68 | 3.04 | 2.33 | 2.84 |
| Os05g0468700 | LOC_Os05g39240 | *OsAMT2;1* | Ammonium transporter. | -2.47 | 1.06 | 1.83 | 1.02 | 0.84 | 2.53 |
| Os05g0474600 | LOC_Os05g39690 |  | Similar to Aldose reductase-related protein. | 1.68 | 0.93 | 2.53 | 2.03 | 1.43 | 1.39 |
| Os05g0537100<sup>3</sup> | LOC_Os05g46020 | *OsWRKY7/10* | WRKY transcription factor 7/10. | -2.12 | 2.25 | 3.13 | 2.15 | 1.83 | 2.35 |
| Os06g0134900<sup>3</sup> | LOC_Os06g04390 |  | Conserved hypothetical protein. | 1.26 | 2.93 | 2.91 | 1.90 | 0.76 | 0.26 |
| Os07g0679300<sup>3</sup> | LOC_Os07g48160 | *α-Gal III* | Similar to Alpha-galactosidase precursor. | 1.60 | 2.40 | 2.84 | 2.33 | 0.95 | 0.47 |
| Os07g0689600<sup>4</sup> | LOC_Os07g48980 | *OsNAS3* | Nicotianamine synthase 3. | -1.45 | -2.00 | -2.81 | -2.89 | -2.54 | -2.19 |
| Os08g0497600 | LOC_Os08g38880 | *SRWD1* | WD40 subfamily protein, Salt stress. | 1.26 | 2.74 | 2.58 | 2.28 | 0.86 | 0.24 |
| Os10g0508000 | LOC_Os10g36390 |  | Similar to C-terminal peptide-binding protein 1. | -1.21 | -0.39 | -1.26 | -1.90 | -2.41 | -1.80 |
| Os11g0181200<sup>3</sup> | LOC_Os11g07911 |  | Hypothetical protein. | 1.74 | 4.37 | 1.92 | 1.70 | 0.14 | 1.12 |
| Os11g0502700<sup>3</sup> | LOC_Os11g30760 |  | Glycosyl transferase. | -1.84 | 1.47 | 3.24 | 2.60 | 1.79 | 1.68 |

<sup>1</sup>, RNA-seq expression data from González-Schain et al. (2016); <sup>2</sup>, Microarray expression data from Zhang et al. (2012); <sup>3,4,5</sup>, Cross-reference studies (Wilkins et al. 2016; Lafarge et al. 2017; Liu et al. 2020) conducted after González-Schain et al. (2016) and Zhang et al. (2012).

**Fig. 4.**
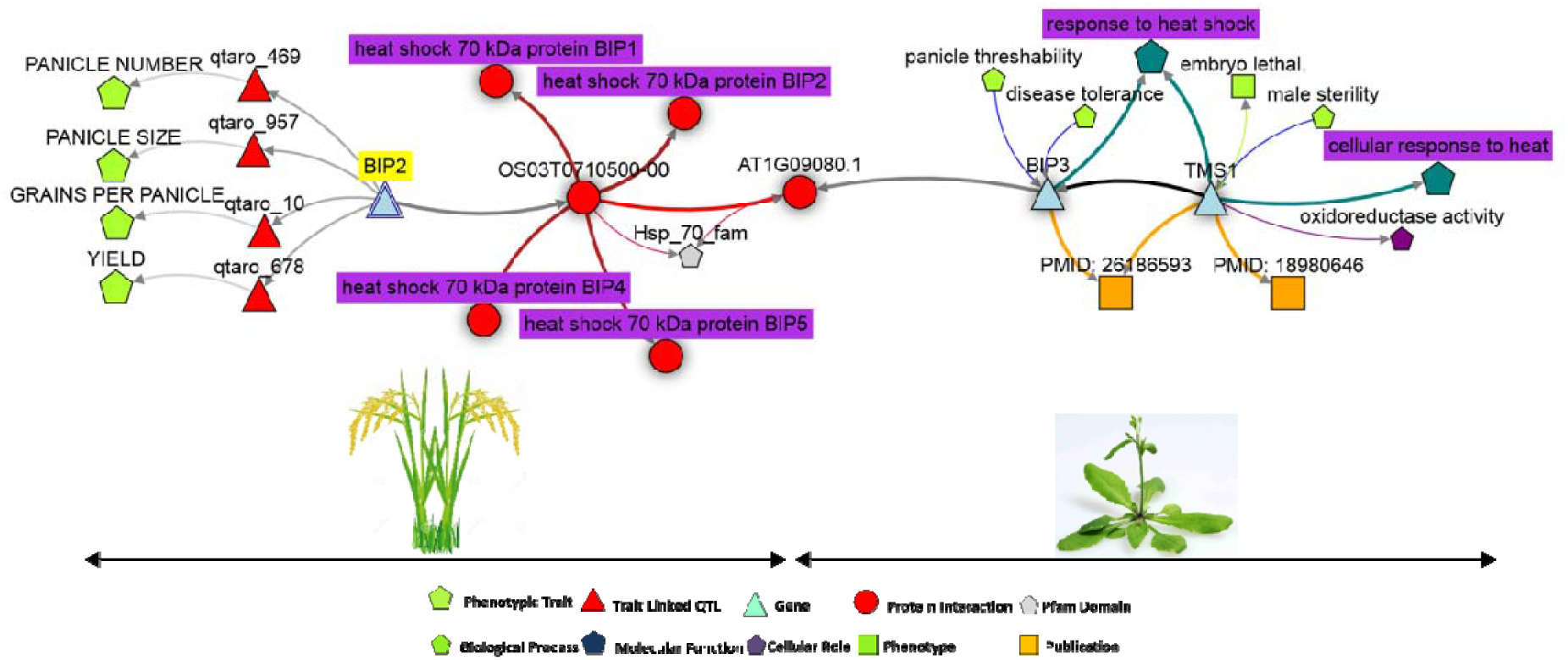
Gene evidence network showing possible relationship between *OsBIP2* and pollen tube thermotolerance. The network was generated with Knetminer by exploring O*sBIP2* (*Os03g0710500*, highlighted in yellow colour) connection with heat tolerance using default parameters. Different colour arrows show link between corresponding entries. This example network provides a proof-of-concept that genes identified in this study are important candidates for reproductive stage heat tolerance.

### CREs in promoters of heat-responsive genes

To investigate whether our core set of heat-responsive genes had common CREs in their promoters, we examined 1.5 kb upstream sequence of all genes and found a total of 27 CREs, which occurred at least once in > 80% of the total genes (**Fig. 5**). Remarkably, the promoters of these genes were enriched with some well-known CREs; such as three heat-responsive (CAATBOX1, HSE, PRECONSCRHSP70A), five dehydration-responsive (ACGTATERD1, MYBCORE, MYB1AT, MYB2CONSENSUSAT, MYCCONSENSUSAT), two sugar-responsive (CGACGOSAMY3, POLASIG1) and three pollen expression specific (GTGANTG10, POLLEN1LELAT52, TAAAGSTKST1) CREs (**Supplementary File-sheet 6**). Furthermore, 17 other stress/stimulus-related CREs including abiotic stress, growth and development, hormone, nutrient and transcription response elements were also found in these genes promoters. Collectively, these results strongly suggest a common regulatory mechanism might exist in rice for mitigating high-temperature stress. In future, it would be interesting to functionally characterize these important genes, along with their promoters, and elucidate their roles in reproductive stage heat-tolerance in rice.

**Fig. 5.**
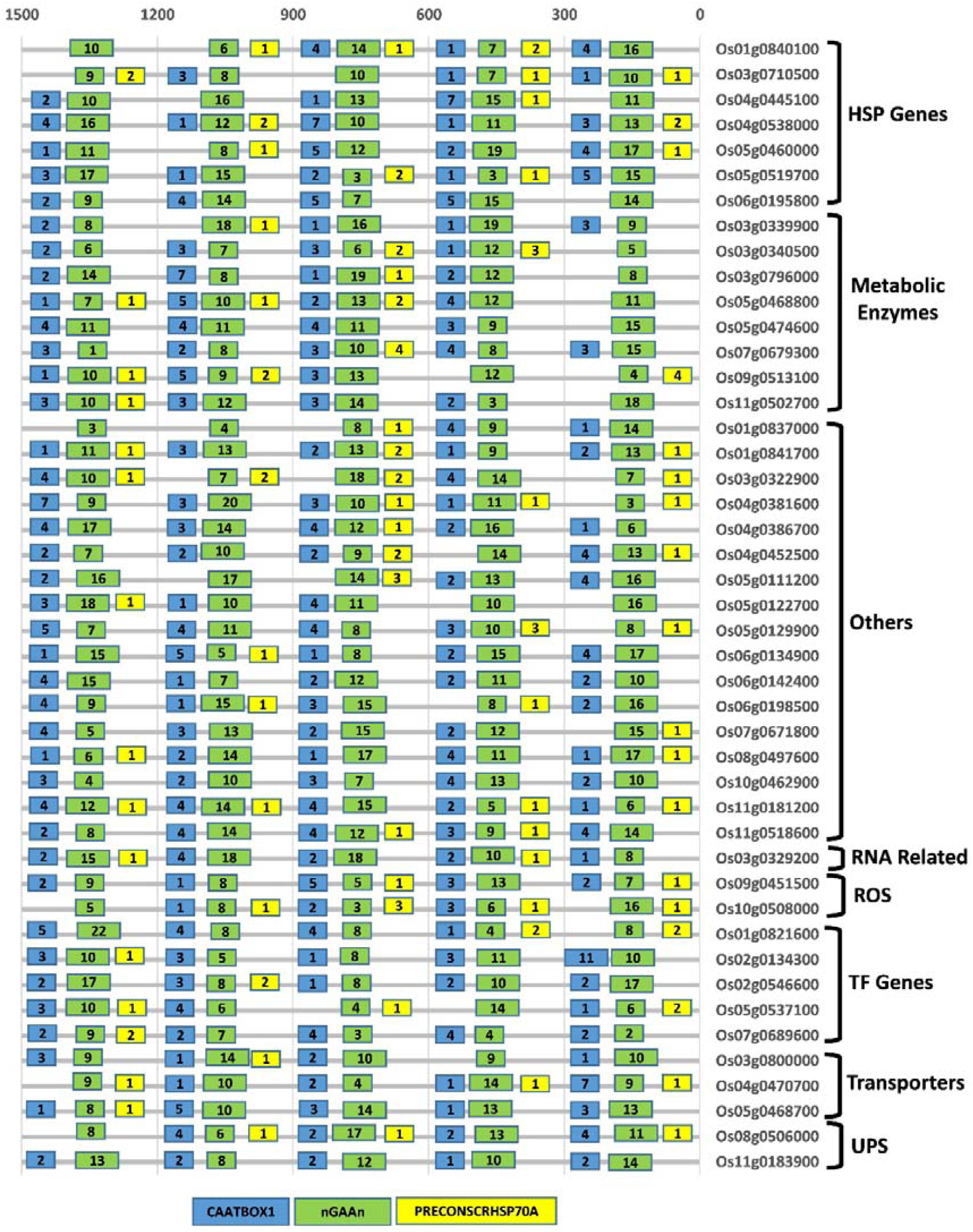
Heat-responsive CREs in core set of heat-responsive genes. Coloured boxes indicate number of CAATBOX1 (blue), nGAAn (green) and PRECONSCRHSP70A (yellow) heat-responsive CREs in 0–300, 301–600, 601–900, 901–1200 and 1201–1500 bp upstream promoter regions. Scale (bp) is presented above the figure. Detailed information of CREs is given in supplementary file-sheet 6.

## Discussion

### Recent advancements in improving reproductive stage heat-tolerance in rice

During recent years, significant progress has been made in screening of heat-tolerant rice germplasm at the reproductive stage and elucidating complex molecular-genetic mechanisms underlying heat-tolerance. For example, Hakata et al. (2017) developed a new assay system for selection of heat-tolerant rice genotypes with no varietal difference in panicle temperature. Lafarge et al. (2017) conducted a genome-wide association analysis on a large set of indica rice germplasm during anthesis and identified several heat stress linked single nucleotide polymorphisms (SNPs) associated with spikelet sterility. Scafaro et al. (2018) demonstrated that a thermotolerant form of photosynthesis-related protein Rubisco activase (Rca) dramatically improves rice growth and grain yield under heat stress conditions. Zhou et al. (2019), by using a hybrid sequencing technology, discovered novel high nighttime temperature stress linked SNPs. Cao et al. (2020) fine mapped a heat tolerance conferring QTL from a wild rice accession and identified two heat-stress induced genes. All these significant advances, along with results presented in this study, could greatly accelerate applied research activities for screening and improvement of heat-tolerance in rice germplasm.

### MQTLs are hotspot genomic regions for development of heat-tolerant rice cultivars

After the advent of genetic markers, mapping of quantitative trait-associated genomic regions in segregating populations has become a routine work. These efforts have yielded in identification of a plethora of trait linked QTLs in diverse genetic backgrounds. Before successfully exploiting these QTLs for trait improvement, identification of consistent QTLs across diverse genetic backgrounds and environments is essential. To achieve this goal, meta-analysis approach has been developed (Goffinet and Gerber 2000) and successfully applied in several agronomically important crops including rice (Raza et al. 2019). In this study, we also performed a meta-analysis on reproductive tissues associated QTLs to identify consistent MQTLs which could be used for improving heat stress tolerance in rice.

To comprehensively identify heat-responsive genomic regions during rice reproductive development, we integrated data of all known heat stress associated QTLs (**Table 1**). According to the definition of meta-analysis, only those genomic regions are considered as MQTLs which harbour at least two original QTLs from different experiments. Our analysis detected 35 MQTLs which contained 2–9 original QTLs (**Fig. 1**). Significant reductions were observed in 95% confidence intervals of 35 MQTLs than the average of their corresponding initial QTLs. Physical and genetic intervals of > 65% MQTLs were narrower than 2 Mb and 5 cM, respectively, and these could be reduced further (**Table 2**). Fine mapping paves the way for identification, cloning and functional characterization of candidate genes underlying the QTL regions (Ye et al. 2015b), which could be further used for trait improvement through MAS. Overall, our results strongly indicate that majority of the MQTLs identified here are hotspots for heat tolerance and should be exploited through marker-assisted breeding programs for development of heat-tolerant rice cultivars.

### Prioritization of candidate genes by integrating genetics and genomics data

QTL mapping establishes relationships between traits of interest and genomic regions. However, a trait linked QTL usually encompasses a vast number of genes and only a few of them are actually casual genes (Bargsten et al. 2014). Identification of trait linked candidate genes has high practical value in marker-assisted breeding. In recent past, transcriptional changes in rice panicles under high-temperature stress conditions were extensively investigated and several heat-responsive genes have been identified (Zhang et al. 2012; González-Schain et al. 2016; Liu et al. 2020). We took advantage of previous work and compared all our genes underlying the MQTL regions with differentially expressed genes of two prominent studies (Zhang et al. 2012; González-Schain et al. 2016). This comparison significantly reduced the number of heat-responsive candidate genes, as we identified only 45 out of total 6785 genes (fold change > 150) (**Fig. 2, Supplementary File-sheets 1–5**). Our results are clearly statistically significant and this comparative approach will greatly help in establishing genotype-to-phenotype relationships, which will ultimately assist in applied research activities.

### General vs. reproductive tissues-specific heat stress responses

Systematic understanding of all type of heat responses is obligatory to improve rice germplasm in view of current global warming scenario. Any change in plant material, stress condition, tissue sample and statistical analysis can affect natural plant responses. In this study, comparisons with most prominent reports in this field, involving a model heat-tolerant rice cultivar N22, revealed some common molecular heat-responses. For example, Zhang et al. (2012) identified 2449 deregulated genes in heat-treated young florets and González-Schain et al. (2016) identified 630 DEGs in heat-stressed mature florets, among them > 6% and nearly 2% were common with our MQTLs underlying genes, respectively (**Supplementary File-sheets 2–4**). The difference in number of common genes between current work and previous studies could be explained by the difference in duration of applied heat stress, floret age, the approach used for studying expression profiles and statistical analyses. Interestingly, at least 45 genes overlapped among three studies and are most important candidates for heat tolerance in rice (**Fig. 2**). Whereas, distinctly shared genes between current work and Zhang et al. (2012), as well as with González-Schain et al. (2016), could be exploited as molecular markers for evaluation of reproductive stage heat tolerance.

In this study, systematic integration of genetics and genomics data provided some novel sources of heat tolerance which could be further explored for elucidating their innate roles in heat stress responses. Interestingly, rice genome holds a small set of basal heat-responsive genes. Twenty-one out of 45 genes takes part in tissue-independent general response to heat stress (**Supplementary File-sheet 5**), as these were also differentially expressed in heat-stressed seedlings (Mittal et al. 2012). Whereas, remaining 24 genes were either missing or responded differently in Mittal et al. (2012) study, indicating a core set of reproductive tissue-specific heat-responsive genes (**Table 3**). This subset includes *HSP*s, *WRKY* TFs, amino acid transporters, sugar metabolism relevant and several other un-classified genes. Functional characterization of these genes could reveal their regulatory roles in heat tolerance during rice reproductive development.

### Reliability of reproductive tissue-specific heat-responsive candidate genes

Next-generation sequencing technologies have triggered an extensive use of microarray and RNA-seq analyses for studying gene expression and identifying trait linked genes (Jain 2012). Heat-responsive candidate genes identified in this study we differentially expressed in both microarray and RNA-seq based investigations after heat stress application and expression results were further validated by RT-qPCR analyses (Zhang et al. 2012; González-Schain et al. 2016). A significant number of these candidate genes also differentially upregulated in reproductive tissues under natural field conditions (Sato et al. 2011a&b) (**Fig. 3**) and take part in rice environmental gene regulatory influence networks (EGRINs) (Wilkins et al. 2016) that function in harsh agricultural environments (**Table 3**). Additionally, spikelet sterility linked SNPs have been identified in two of our candidate genes (*OsWRKY21* and *OsNAS3*) (Lafarge et al. 2017). Collectively, deregulation of all these candidate genes under normal field conditions, as well as after heat stress application, and enclosure of majority of genes in cross-reference studies endorse their inherent connection with reproductive stage heat tolerance.

### *OsBIP2* and thermotolerance of pollen tubes

To show a possible relationship between *OsBIP2* and thermotolerance of pollen tubes, we generated a gene evidence network based on functional data from *Arabidospsis* genes (**Fig. 4**). Gene-evidence-network showed that *OsBIP2* (a highly expressed gene in reproductive tissues of rice) co-locate with panicle number, panicle size, grains per panicle and yield QTLs (qtaro_10, 469, 678 and 957;) on chromosome 3 and encodes a heat shock 70 domain-containing protein (PF00012). Its orthologs in *Arabidopsis* (*BIP3, TMS1*) play important roles in thermotolerance of pollen tubes under high-temperature conditions (Ma et al. 2015; PMID 26186593). Knockout mutants of *TMS1* display retarded pollen tube growth and reduced fertility under normal conditions (Yang et al. 2009; PMID 18980646) (**Fig. 4**). This example network provides strong evidence that *OsBIP2* have potential function in pollen tubes thermotolerance and genes identified in this study are important candidates for reproductive stage heat tolerance.

### Common regulatory mechanism might exist in rice for heat-tolerance

CREs function as important molecular switches for transcriptional regulation of complex gene regulatory networks during harsh environments. Different signaling pathways regulate stress-responsive genes by cross-talking with CREs present in their promoters (Yamaguchi-Shinozaki and Shinozaki 2005). In this study, the promoters of our core set of heat-responsive genes were enriched with some functionally characterized reproductive tissue-specific and heat stress associated CREs (**Fig. 5**). For example, heat shock factors bind to heat shock element (HSE; nGAAn) containing gene promoters and trigger heat stress response (Sakurai and Enoki, 2010). CAAT-box and PRECONSCRHSP70A elements have quantitative effects on expression of heat-shock genes in transgenic tobacco and *Chlamydomonas*, respectively (Rieping and Schöffl 1992; von Gromoff et al. 2006). GTGANTG10, POLLEN1LELAT52 and TAAAGSTKST1 are pollen expression specific CREs in rice (Sharma et al. 2011). CGACGOSAMY3 and POLASIG1 are α-amylase genes specific CREs and play roles in sugar production from stored starch (Hwang et al. 1998; O’Neill et al. 1990). Similarly, ACGTATERD1, MYBCORE, MYB1AT, MYB2CONSENSUSAT and MYCCONSENSUSAT are dehydration-responsive CREs found in the promoter of *Arabidopsis* universal stress gene (*AtUSP*) (Bhuria et al. 2016). In our core set of genes, 25–76 copies of HSE (nGAAn) and at least one copy of other above mentioned CREs is present in > 80% genes (**Supplementary File-sheet 6**). This single copy is sufficient to induce a heat stress response, as Narusaka et al. (2003) reported that a single copy of drought-responsive element (DRE) is sufficient for drought-responsive gene expression. Enrichment of our core set of genes promoters with reproductive tissue-specific and heat stress associated CREs strongly suggest a common regulatory mechanism might exist in rice for heat-tolerance during reproductive development.

## Conclusion

Development of heat-tolerant cultivars is a top priority in current global climate change scenario. Marker-assisted breeding can greatly accelerate applied research activities by introgressing stable large phenotypic effect QTLs/genes in advance breeding material. MQTLs identified in this study are most reliable genomic regions for marker-assisted introgression and exploration of novel heat-tolerance candidate genes. Similarly, candidate genes identified from MQTLs showed a significant response to heat stress and are important candidates for improvement of heat tolerance in rice. Moreover, gene evidence network between *OsBIP2* and thermotolerance of pollen tubes also highlights potential benefits of data integration in establishing genotype-to-phenotype relationships. Finally, enrichment of candidate genes promoters with heat stress associated CREs indicate some common regulatory mechanism exists in rice and provides new insights into the molecular basis of heat tolerance. To the best of our knowledge, this is the first report on meta-analysis of reproductive stage heat stress associated QTLs in rice and can improve the success of future rice breeding efforts.

## Authors’ contributions

Q.R and A.R conceived the idea. A.R retrieved QTL data, genes information & expression data. Q.R conducted meta-analysis, drew figures and drafted the manuscript. K.B and M.S reviewed and edited the manuscript. M.S planned the projects, acquired funding and coordinated the collaboration between project partners. All authors listed have made direct and substantial efforts for improving the manuscript and approved the final version.

## Appendix A. supplementary data

All supplementary data is attached with this article in the form of single Microsoft Excel file.

## Code availability

Not applicable

